# Sterol-O acyltransferase 1 is inhibited by gga-miR-181a-5p and gga-miR-429-3p through the TGFβ pathway in endodermal epithelial cells of Japanese quail

**DOI:** 10.1101/569806

**Authors:** Han-Jen Lin, Chiao-Wei Lin, Harry J. Mersmann, Shih-Torng Ding

**Author notes:** 1st set of equal contributors. 2nd set of equal contributors, these authors also contributed equally to this work (Chiao-Wei Lin, Harry J. Mersmann). **Corresponding author:** Dr. Shih-Torng Ding.

## Abstract

Nutrients are utilized and re-constructed by endodermal epithelial cells (EECs) in yolk sac membranes in avian species. Sterol-O acyltransferase 1 (SOAT1) is the key enzyme to convert cholesterol to cholesteryl ester for delivery to growing embryos. During development, absorption of yolk is matched with significant changes of SOAT1 mRNA and enzyme activity. miRNAs regulate angiogenesis and metabolism during mammalian development. However, the involvement of miRNAs in lipid utilization during avian embryogenesis remains ambiguous.

Using a miRNA sequencing technique, we found several candidate miRNAs and confirmed expression patterns with real time PCR. They were selected for as candidates targeting the receptor (TGFβ receptor type 1, TGFBR1) that may regulate SOAT1. Similar to SOAT1 mRNA accumulation, the gga-miR-181a-5p expression was gradually elevated during development, but the concentration of gga-miR-429-3p was in the opposite direction. Transfection with gga-miR-181a-5p or gga-miR-429-3p inhibited TGFBR1 and SOAT1 in EECs. The 3’ untranslated region (3’UTR) of TGFBR1 was then confirmed to be one of the targets of gga-miR-181a-5p and gga-miR-429-3p. Taken together, expression of miRNAs during embryonic development regulates SOAT1 expression by inhibiting the 3’UTR of TGFBR1. This is indicative of possible regulation of avian yolk lipid utilization and modification of hatchability by changing miRNA expressions.

## Introduction

SOAT1 (sterol-O acyltransferase 1), also named ACAT1 (acyl-Coenzyme A: cholesterol acyltransferase 1), is the key enzyme to catalyze cholesterol conversion into cholesteryl ester, by adding fatty acyl coenzyme A; thus, a less polar molecule is produced [1]. Yolk sac membrane (YSM), a three-layer extraembryonic tissue, serves crucial roles for avian embryos during the entirety of embryonic development. We demonstrated that SOAT1 activity in endodermal epithelial cells (EECs, the third layer of YSM) was activated by specific nutrients and hormones through the cAMP-dependent PKA signaling pathway, and accumulated more cholesterol ester in EECs [2].

The diversity of bio-functions and involvement of non-coding RNAs has raised considerable issues. Non-coding RNAs include short (microRNAs, miRNAs) and long non-coding (lncRNAs), ribosomal (rRNAs), transfer (tRNAs), small nuclear (snRNAs), small nucleolar (snoRNAs), transfer-messenger (tmRNAs) and telomerase RNAs [3]. The functions and regulations of miRNAs have been examined in mammalian species for decades. Mainly, mature miRNAs are paired to 3’ untranslated regions (UTR) or 5’UTR by identifying seed regions of target genes [4].

During avian embryonic development, the comprehensive whole mount in situ hybridization expression analysis of 111 mature miRNA sequences in chicken embryos revealed that miRNAs showed a variety of patterns in the early stages of development [5]. Tissue specific-expressed miRNAs were also found to regulate lipid metabolism and cell proliferation at later stages in chicken embryonic livers [6]. Some miRNAs were extracted and detected in albumen and yolk from chicken unembryonated eggs. This suggested that miRNA transport from laying hens into albumen or yolk would be efficient to facilitate normal embryonic development by continually supplying miRNAs to growing embryos during nutrient uptake [7]. Nutrient absorption and reassembly in YSM has been confirmed [2, 8]. However, the miRNA expression patterns of YSM and the crucial linkages between embryos and yolk remain unclear during development.

The TGFβ signaling pathway is substantial in development [9]. The TGFβ family is involved in paracrine signaling and can be found in different tissue types, including brain, heart, kidney, liver, and sex organs. TGFβ receptor types I and II have similar ligand-binding affinities and can only be distinguished by peptide mapping. Both receptor type I and II have high affinities for TGFβ1, but low affinities with TGFβ2. Overall activation of the TGFβ signaling pathway is through TGF family-ligand binding followed by continuous phosphorylation of the type I and then type II receptor. The Smad2/3 proteins, known as signal transmitters, are phosphorylated after TGFβ receptor activation. Smad4 then joins with Smad2/3 to form the transcription factor complex to enter the nucleus and regulate promoter regions of target genes.

The relationships between SOAT1 and the TGFβ signaling pathway during lipid metabolism are less discussed. Although TGFβ alters cellular cholesterol metabolism in smooth muscle cells by increasing LDL receptor expression and simulating substrate binding (LDL), as well as enhancing delivery of cholesterol, the SOAT1 activity is not changed [10]. Similarly, TGFβ increases cholesterol efflux in macrophage-derived foam cells, but the SOAT1 mRNA expression (analyzed by Northern blotting) remains unchanged after TGFβ stimulation [11]. However, exogenous TGFβ1 upregulates SOAT1 expression and activity during transition of human monocytes into macrophages [12]. TGFBR1 proteins are detected from early stages in chicken embryos [13]. Although the Smad3 transcription factor binding region in the SOAT1 promoter is predicted by Genomatix, the detailed mechanism of the TGFβ signaling pathway regulating SOAT1 needs to be clarified.

In this study, we demonstrated the miRNA profiling and miRNA-mRNA interaction in primary EECs from YSM of Japanese quail. The aim of the current research was to discover potential miRNAs involved in the TGFβ signaling pathway and modulation of SOAT1 expression in EECs during embryonic development.

## Material and methods

All animal studies were approved by the Institutional Animal Care and Use Committee (IACUC) of the National Taiwan University. The IACUC Approval No: NTU107-EL-00148.

### microRNA sequencing

The microRNA sequencing of yolk sac membranes (YSM) during Japanese quail (*Coturnix coturnix*) embryonic development was analyzed by PhalanxBio Inc. (Hsinchu, Taiwan). For a better understanding of the overall miRNA expression profiles, samples of YSM were collected at embryonic day 5 (ED5), ED10, ED15, and post-hatch day 2 (PH2); one sample was used at each time point. Total RNA was sequenced by Illumina HiSeq2500; raw data was compared with references to a chicken microRNA database, miRBase v21, for comparison of miRNA precursors and mature miRNA sequences.

### Prediction of microRNA targeting genes

Two software programs were applied to predict the unknown chicken miRNAs targeting SOAT1 and the potential targets of selected miRNAs. We searched for miRNA candidates that affect SOAT1 and the transforming growth factor-beta signaling pathway (TGFβ signaling pathway) using miRDB (http://www.mirdb.org/miRDB/index.html) [14, 15] and TargetScanHuman 7.2 (http://www.targetscan.org/vert_72/) [16]. We confirmed the selected miRNAs with miRNA sequencing data. The sequences of selected miRNA are listed as Table 1. SMAD3 (SMAD family member 3, a family of proteins similar to the Drosophila gene ‘mothers against decapentaplegic’ (Mad) and the C. elegans gene Sma), was one of the main signal transducers in the TGFβ signaling pathway for the SMADs complex assembly and entrance into the nucleus. According to results of Genomatix prediction, the SMAD3 transcription factor binding site was present in the SOAT1 promoter. Therefore, we hypothesized that the miRNAs affect both SOAT1 and factors in the TGFβ signaling pathway.

**Table 1.**
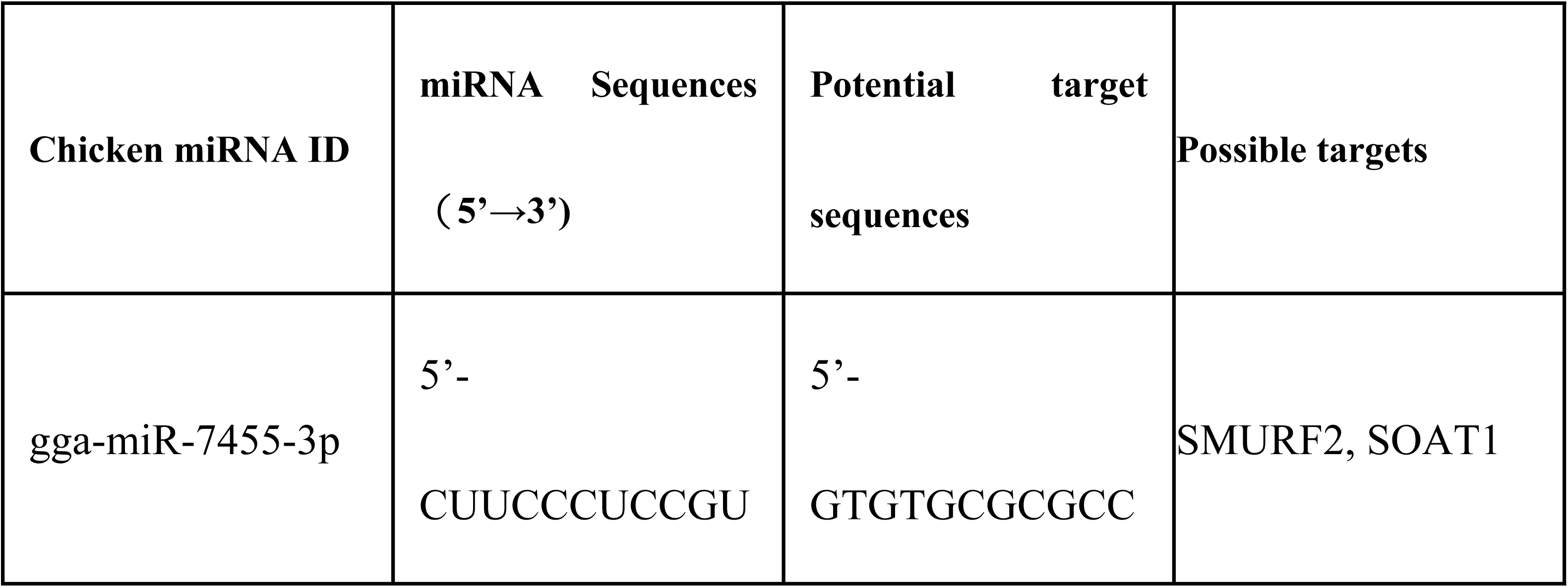
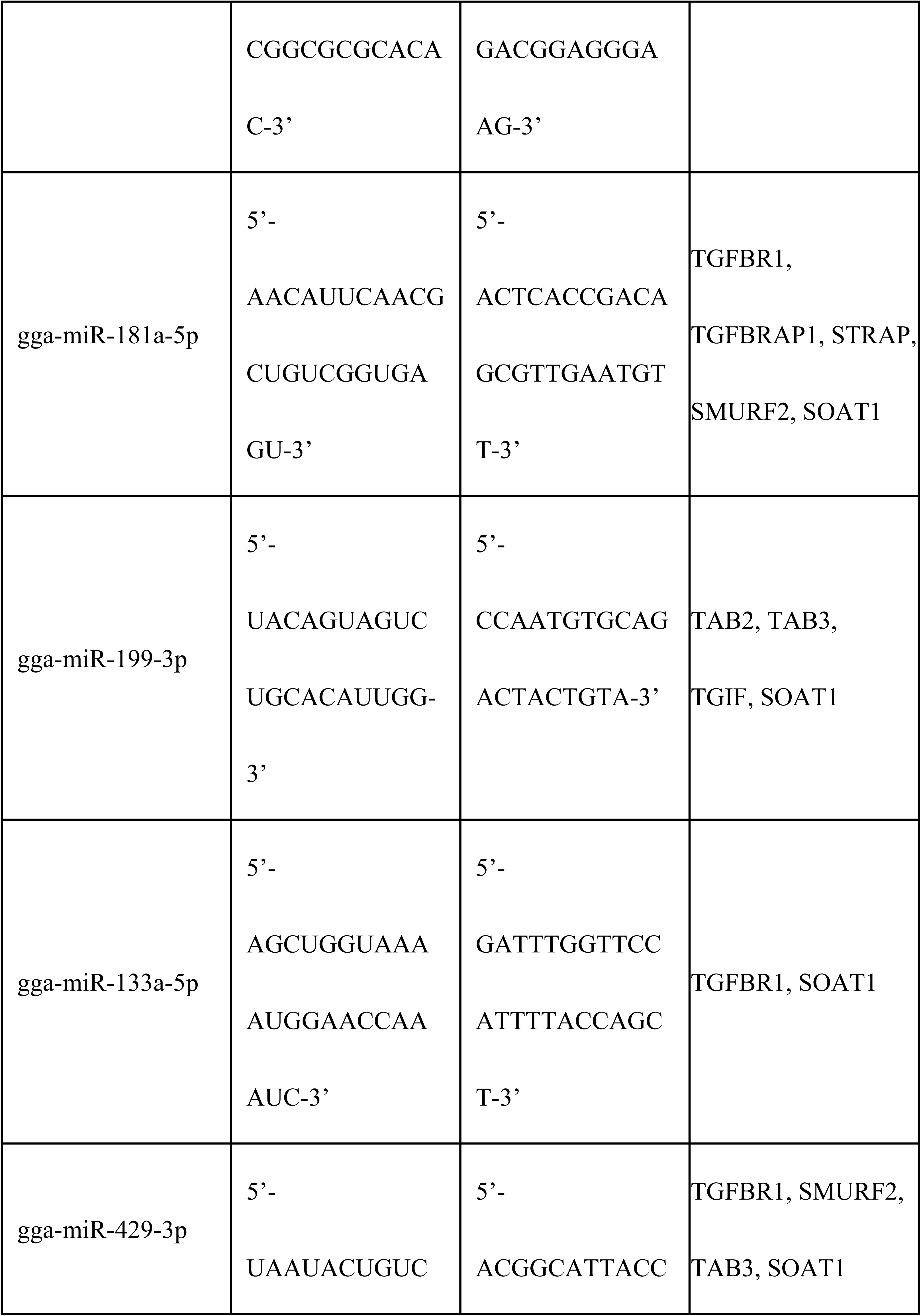
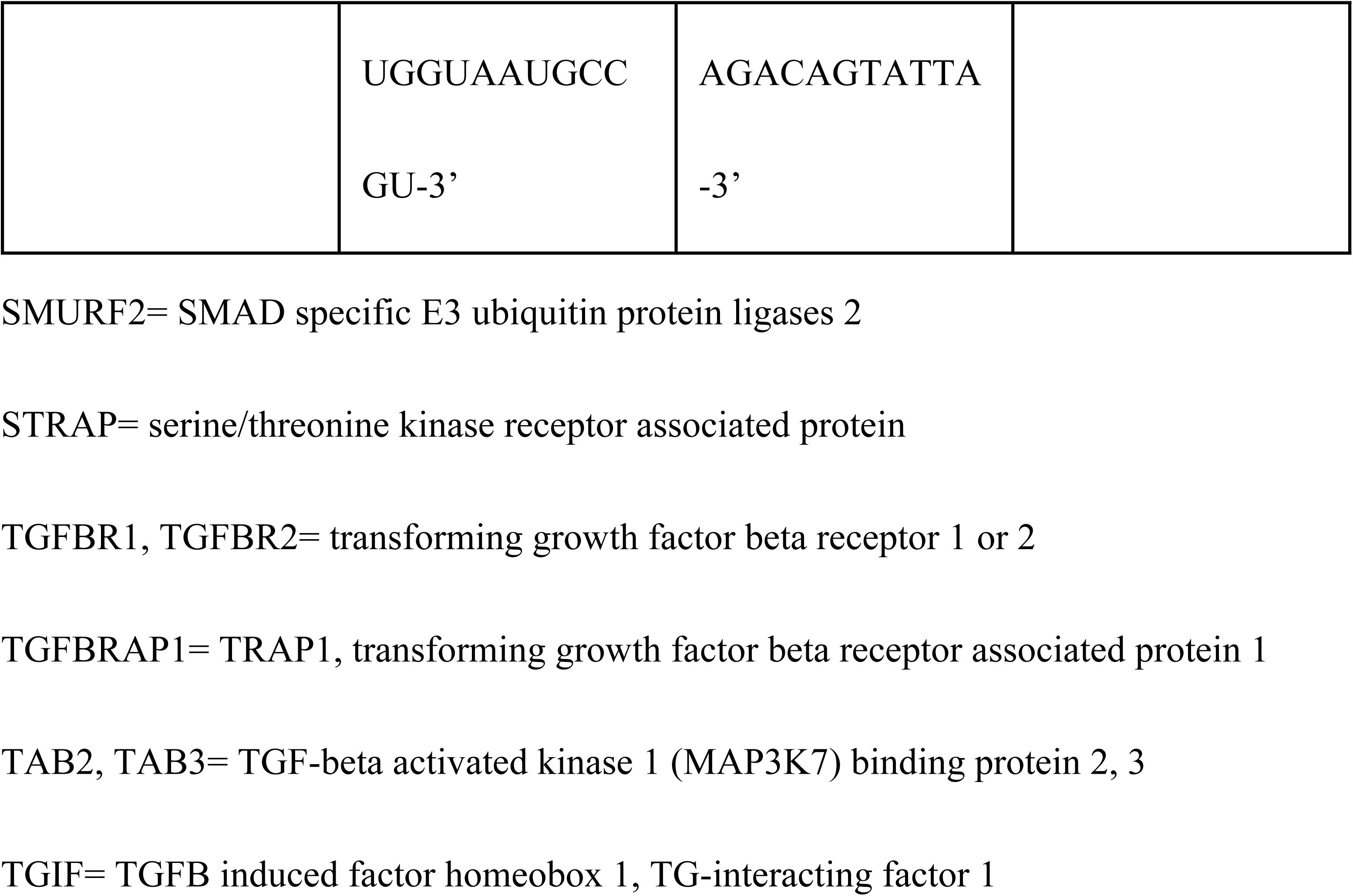
The list of selected miRNAs.

### Validation of microRNA expressions in YSM tissues of Japanese quail

Total RNA of YSM tissues from four embryonic days were extracted by GENEzol^™^ Reagent (New Taipei City, Taiwan). The miRNAs were modified by polyadenylation at the 3’ end and then reverse transcribed into the cDNA of miRNA using the miScript PCR Starter Kit (#218193, Qiagen, Valencia, CA, USA) with an oligo dT primer (with a universal tag). The custom miScript Primer Assays (as forward primer, Table 2) were designed to identify different miRNAs and miScript Universal Primer was used as reverse primer. Real-time PCRs were analyzed by SensiFAST™ SYBR^®^ Hi-ROX Kit (BIO-92020, Bioline, London, UK). A PCR program was used as described: 15 minutes at 95°C, 40 cycles of 15 seconds at 94°C for denaturation, 30 seconds at 55°C for primer annealing, 30 seconds at 70°C for extension, and 1 minute at 70°C for final extension. All kits and primer assays were purchased from commercial sources and were used according to manufacturer instructions here and elsewhere in this manuscript.

**Table 2.**
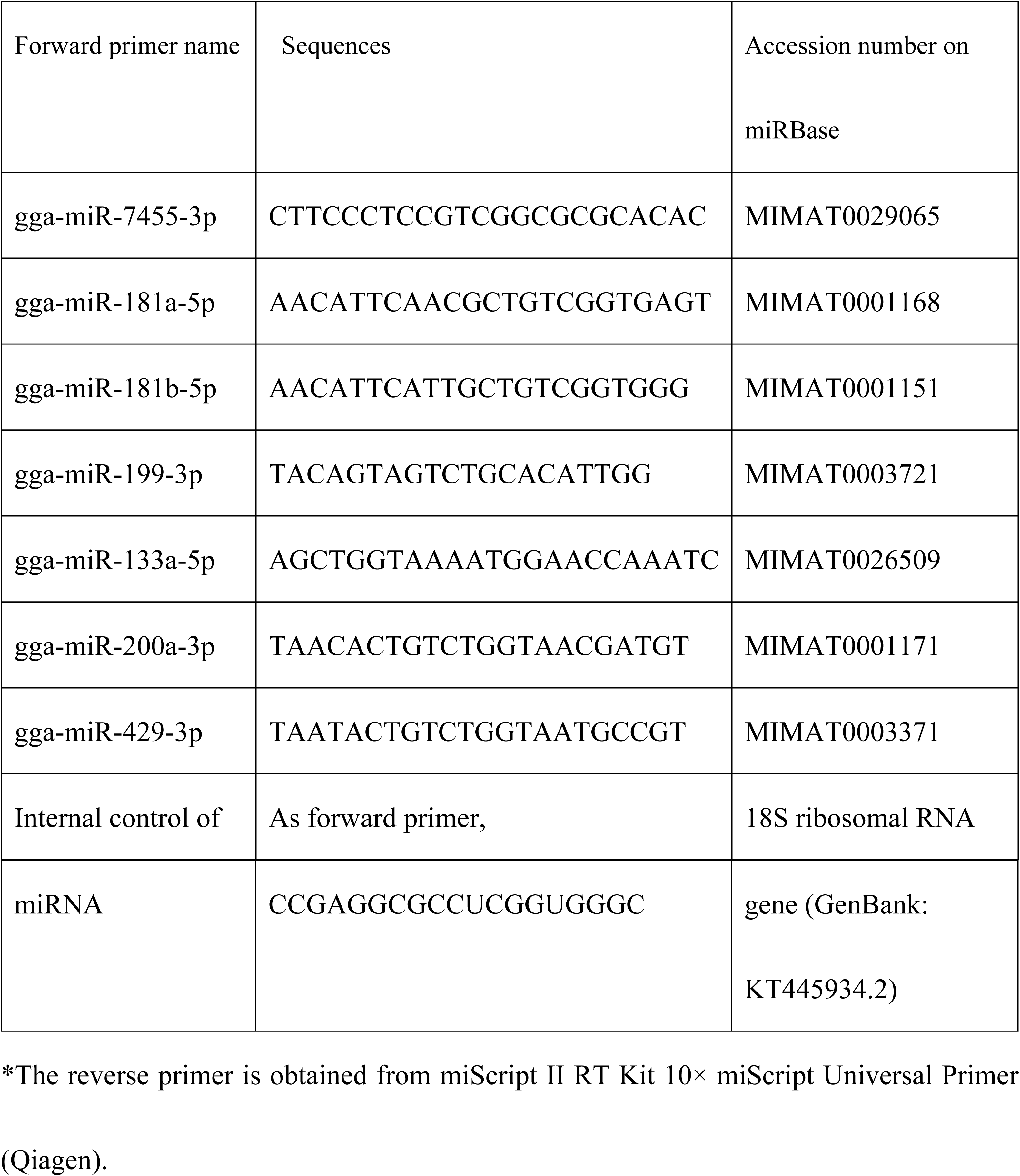
The miScript primer list.

### Cell culture system

Isolation of endodermal epithelial cells (EECs) and the culture system were modified from the published procedure [8]. In short, YSM tissues from day 5 embryos were treated with collagenase (collagenase type 4, 17104019, ThermoFisher, Waltham, MA, USA) to partially digest the extracellular matrix and facilitate cell isolation [17]. We collected six YSM (from six embryonic day five embryos) to isolate EECs; these were pooled as one sample for the experiment. EECs were cultured in DMEM/ F12 (pH 7.4, 12400–024, ThermoFisher) with 10% new born calf serum (16010–159, ThermoFisher) and 1% Penicillin-Streptomycin-Amphotericin B Solution (PSA, 03-033-1B, Biological Industries, Cromwell, CT, USA).

To emphasize functional effects, selected miRNAs were transient transfected into EECs after seeding for 48 hours. The culture medium was changed before transfection. The transfection complexes were prepared with 5 nM miRNA mimics or a 5 nM siRNA negative control (AllStars Negative Control siRNA, 5’-UUCUCCGAACGUGUCACGU-3’) in DMEM/F12 using 3 μL HiPerFect^®^ Transfection Reagent. The custom miScript miRNA mimics, negative control, and HiPerFect^®^ Transfection Reagent were purchased from a commercial source (Qiagen).

The HEK293T cell line was used for validation of the target pairing between miRNAs and target sequences by the luciferase reporter assay. The 293T cells were cultured in DMEM (pH 7.4, 12800-017, ThermoFisher) with 10% fetal bovine serum (SH30071.02, GE Healthcare Life Sciences, Utah, USA) and 1% PSA.

### Real time PCR for measuring gene mRNA accumulations

The total RNA of YSM tissues or EECs was extracted using the GENEzol^™^ Reagent (New Taipei City, Taiwan), followed by reverse transcription with a High Capacity cDNA Reverse Transcription Kit (4368814, ThermoFisher). The cDNA was stored at −20°C. The specific primers for quail gene expressions were designed by Primer3 (http://frodo.wi.mit.edu/primer3/) and listed below (Table 3). The reactions were prepared using a SensiFAST^™^ SYBR^®^ Hi-ROX Kit and 0.3 μM specific primers. The program used was: 3 minutes at 95°C, 40 cycles of 5 seconds at 95°C and 30 seconds at 60°C for annealing, with final extension for 1 minute at 60°C.

**Table 3.**
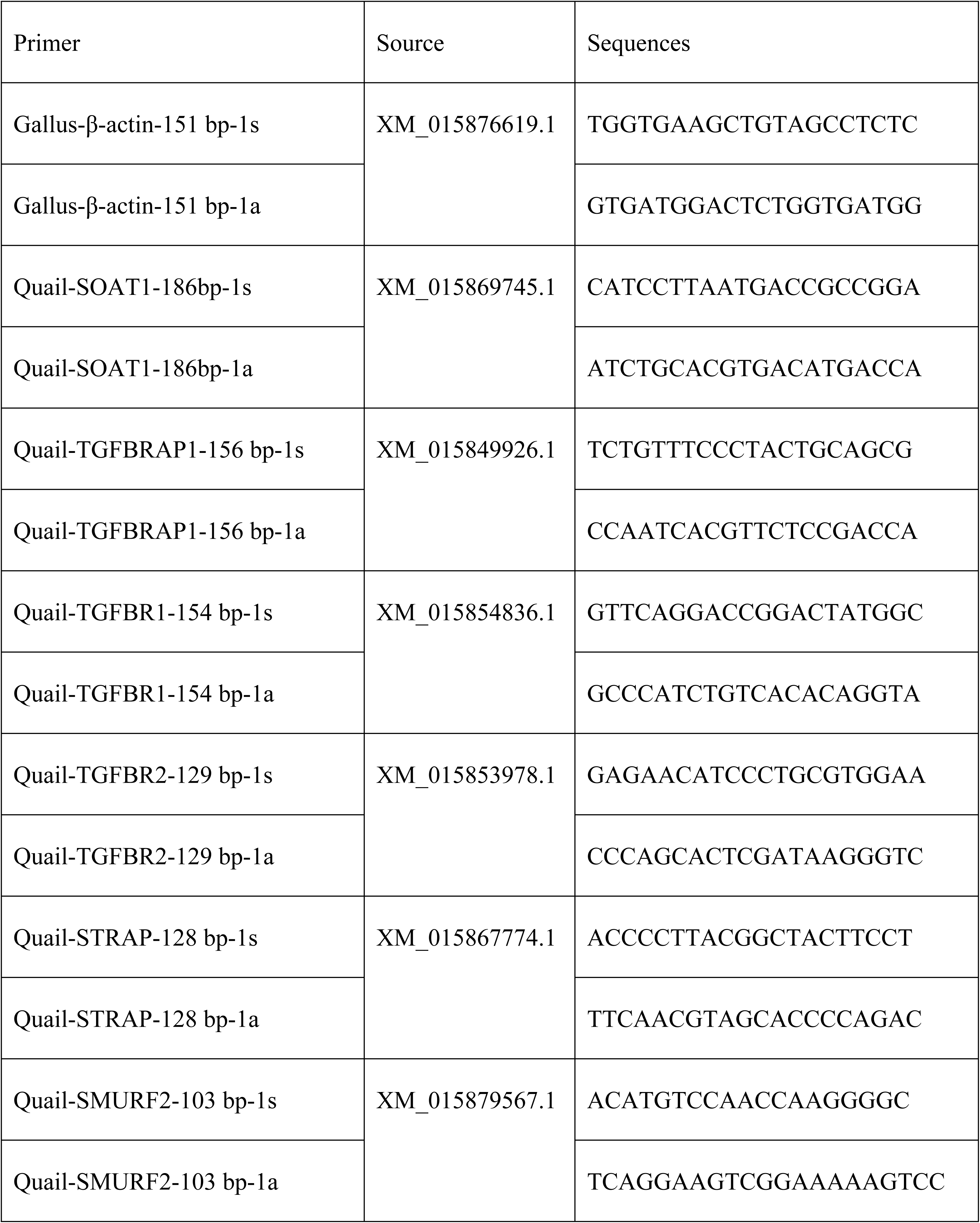
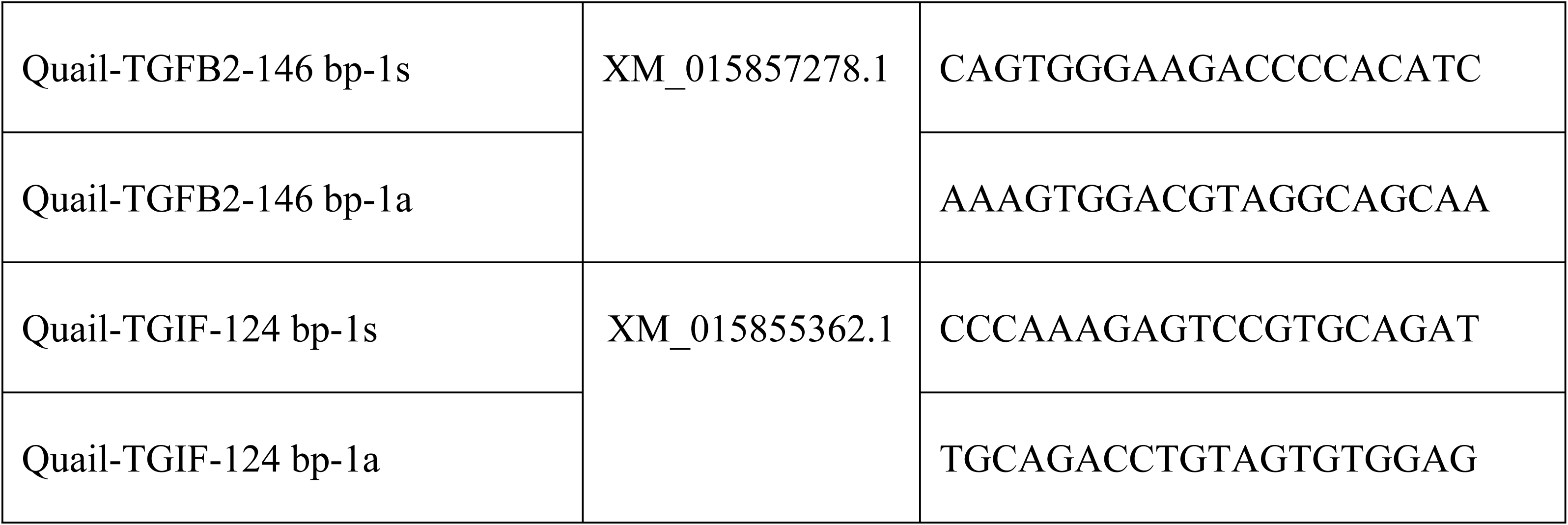
The Real time-PCR primers

### Luciferase plasmid construction and luciferase reporter assay

To verify the miRNA-mRNA pairing between miRNAs and 3’UTR of chicken transforming growth factor beta receptor 1 (TGFBR1, NM_204246.1), the synthetic WT sequences of the 3’UTR (Genomics, New Taipei City, Taiwan) were amplified and cloned into a pmirGLO Dual-Luciferase miRNA Target Expression Vector (E1330, Promega, Madison, WI, USA) at SacI and XhoI restriction sites. The primers used for amplifying the mutated sequence are listed as Table 4. The 3’ UTR of TGFBR1 was predicted to contain two gga-miR-181a-5p binding sites and 1 gga-miR-429-3p binding site. Therefore, the synthetic mutants of TGFBR1 3’UTR were separately inserted into pmirGLO vectors. The pmirGLO-mutant-3’UTR plasmids, which had two 7 bp substitutions in the seeding regions of miR-181a-5p (MUs: TTGAATG→ GGTCCGT), and one 7 bp substitution in the seeding region of miR-429-3p (MU: TAATACT→ GCGCGCG) were all sequenced.

**Table 4.**
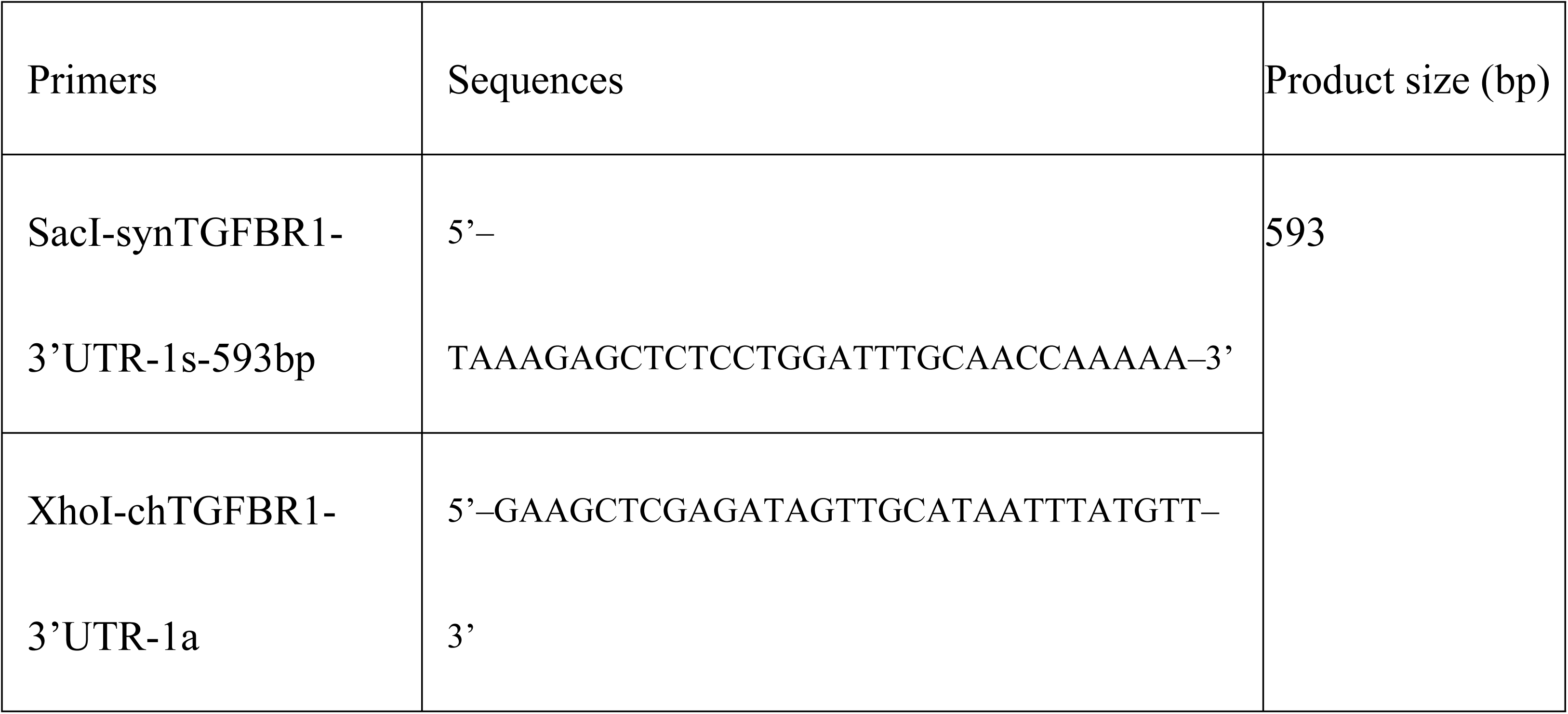
Primers for synthetic 3’UTR amplification.

The 293T cells at a density of 3 × 10^4^ cells/ well on 96-well plates were cultured in DMEM medium with 10% fetal bovine serum and 1% PSA. When the cells reached 60% to 70% confluence, pmirGLO-syn-3′UTR (100 ng) or pmirGLO-syn-mu-3’UTR-181a-5ps (100 ng) or pmirGLO-syn-mu-3’UTR-429-3p (100 ng) were co-transfected with a negative siRNA control or 5 to 15 nM gga-miR-181a-5p, gga-miR-199-3p, gga-133a-5p, or gga-miR-429-3p mimics (Qiagen) using 0.65 μL of PolyJet^™^ (SL100688, SignaGen^®^ Laboratories, Rockville, MD, USA). The luciferase expression was measured by the Dual-Glo Luciferase Assay System (Promega) after transfections for 24 hours, and was detected by SpectraMax i3 and SoftMax Pro 7.0 (Molecular Devices, San Jose, CA, USA).

### SDS-polyacrylamide gel electrophoresis and immunoblotting

EECs were transfected by GenMute™ siRNA Transfection Reagent (SL100568, SignaGen^®^ Labroatories) after culturing for 48 hours. Total protein of EECs was extracted with 1X RIPA buffer (20–188, Merck, Darmstadt, Germany), supplemented with Halt^™^ Proteinase & Phosphatase Single-Use inhibitor cocktail (78442, ThermoFisher). Total proteins were collected by and centrifugation procedure (17000 g at 4 °C for 30 minutes) to remove mitochondria, cell membranes, nucleus and others. The supernate was stored in - 80 °C for Western blotting following the previously described procedure (2). In brief, 15 μg protein/ per sample as determined using the Pierce^™^ BCA Protein Assay kit (23227, ThermoFisher) were subjected to 10% SDS-PAGE gel with 80V for 130 minutes, and the separated proteins were electrophoretically transferred to a polyvinylidene difluoride (PVDF) membranes (NEF1002001PK, PerkinElmer, Waltham, MA, USA) by 400 mA for 75 minutes. Nonspecific binding sites were blocked with 5% skim milk for 1 hour at room temperature. SOAT1 was detected with rabbit anti-mouse SOAT1 primary antibody (antibody diluted 1:300, orb100781, Biorbyt, Cambridgeshire, UK) followed by incubation with anti-rabbit IgG HRP-linked secondary antibody (1:5000, 7074S, Cell Signaling, Danvers, MA, USA). TGFBR1 was detected with rat anti-mouse TGFBR1 primary antibody (1:300, sc-101574, Santa Cruz, Dallas, Texas, USA) and followed by incubation with anti-rat IgG HRP-linked secondary antibody (1:5000, bs-0293G-HRP, Bioss, Woburn, MA, USA). The β-actin protein (1:1000, sc-4778, Santa Cruz) was detected as an internal control. The target proteins were detected with the Clarity^™^ Western ECL Blotting Substrate (#170-5061, Bio-Rad, Hercules, CA, USA). The sizes of proteins were estimated with a PageRuler^™^ Prestained Protein Ladder (10-180 kDa) (26616LCS, ThermoFisher). Protein quantifications were performed with Bio-Rad ChemiDoc Touch Imaging program.

## Statistical analysis

All data were analyzed by one-way analysis of variance. The major effect between treatments was determined by Dunnett’s multiple comparison post-hoc test. The significance level used was at P≤ 0.05.

## Results

### The discovery of candidate miRNAs involving in SOAT1 regulation during embryonic development

The aim was to find potential miRNAs for direct or indirect modulation of SOAT1 expression. The miRNA database for Japanese quail was not yet available, therefore, we used the database from chickens (*Gallus gallus*). The miRNA lengths were mostly concentrated at 22 bps (Fig 1A), and the clustering analysis showed (Fig 1B) that there were 30 miRNAs with the most variance among the four developmental time points. To clarify the regulation of SOAT1, two online searching tools, miRDB (http://www.mirdb.org/miRDB/) and TargetScanHuman 7.2 (http://www.targetscan.org/vert_72/) were used to find potential miRNAs targeting SOAT1. These predictions were further confirmed by miRNA sequencing data.

**Fig 1.**
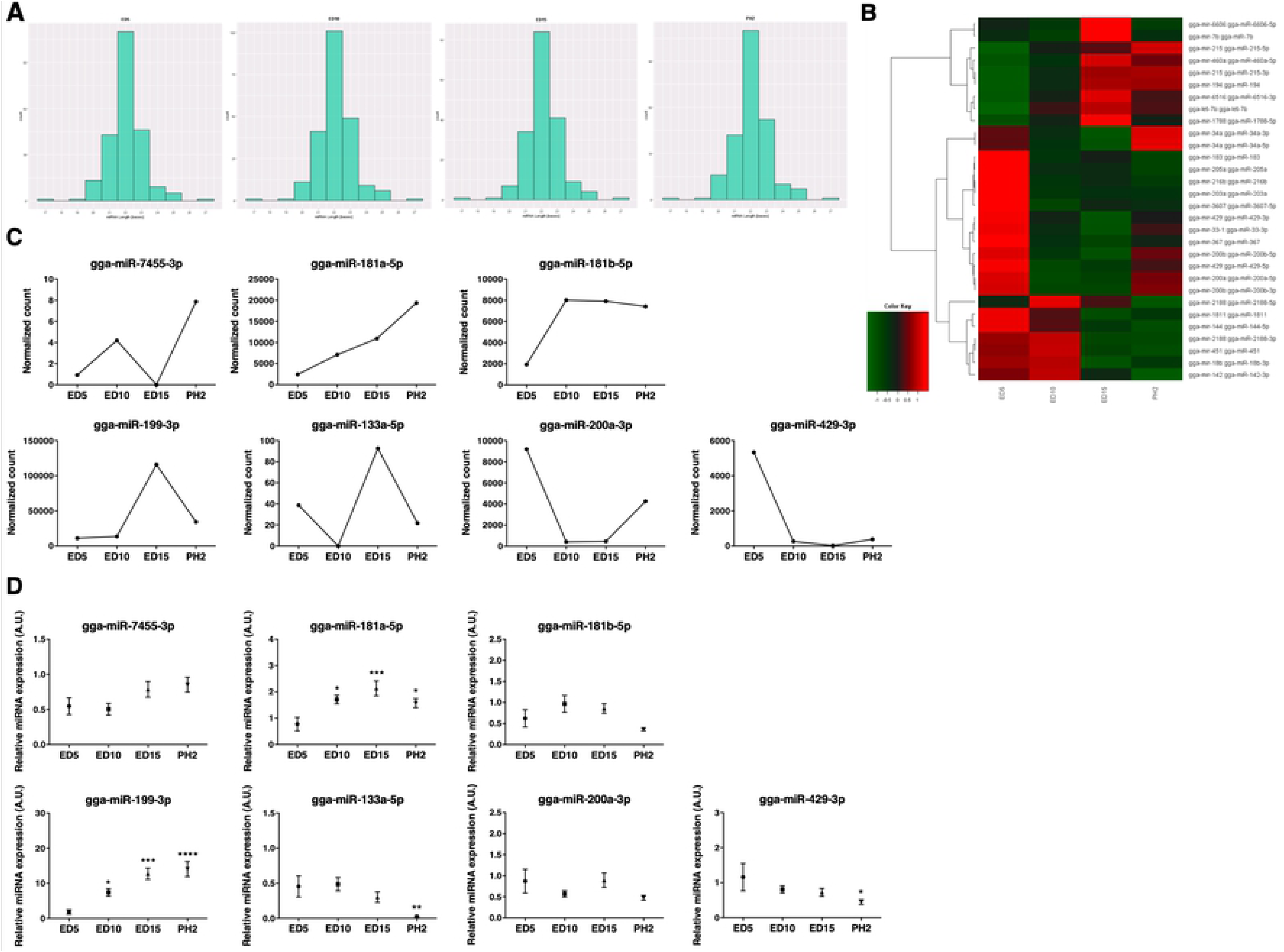
Specific miRNA expressions during embryonic development of Japanese quail. Embryonic day was as ED, and post-hatch day was as PH. (A) Base pair length of miRNAs; the most frequent appearance of miRNAs was at 22 bp. (B) Cluster analysis of miRNA sequencing. Clustering was performed to visualize the correlations among the replicates and varying sample conditions. A subset of microRNAs that exhibited the most variance was selected for cluster analysis. The number of microRNAs clustered was 250. (C) The sequencing data set for gga-miR-7455-3p, gga-miR-181a-5p, gga-miR-181b-5p, gga-miR-199-3p, gga-miR-133a-5p, gga-miR-200a-3p, and gga-miR-429-3p. One sample was used at every time-point in the sequencing result. (D) Confirmation of miRNA expressions by real-time PCR. Every point included seven to nine samples per group. Data were expressed as mean ± S.E.M. Statistical significance was determined by one-way analysis of variance. Dunnett’s multiple comparison test was used to evaluate differences between means. A significant difference from ED5 samples was indicated as*P≤0.05, **P≤0.01, ***P≤0.005 or ****P≤0.001.

Seven miRNAs were selected and listed according to the scoring by miRDB (Table 1). The higher scores indicated the more confidence in prediction algorithm. They also showed expression patterns according to miRNA sequencing (Fig 1C). Expressions of miRNAs were further verified by real-time PCR on YSM samples (Fig 1D). The seven miRNAs were gga-miR-7455-3p (MIMAT0029065; prediction score-94), gga-miR-181a-5p (MIMAT0001168; score-88), gga-miR-181b-5p (MIMAT0001151; score-88), gga-miR-199-3p (MIMAT0003721; score-81), gga-miR-133a-5p (MIMAT0026509; score-80), gga-miR-200a-3p (MIMAT0001171; score-73), and gga-miR-429-3p (MIMAT0003371; score-71).

Because we only collected and analyzed one sample at each timepoint to sequence, the correlations between miRNA sequencing and real-time PCR were calculated. The results showed that the most similarity between the 7 miRNAs was with gga-miR-429-3p (about 83%), gga-miR-199-3p (61%), and gga-miR-181a-5p (50%) (Fig 2).

**Fig 2.**
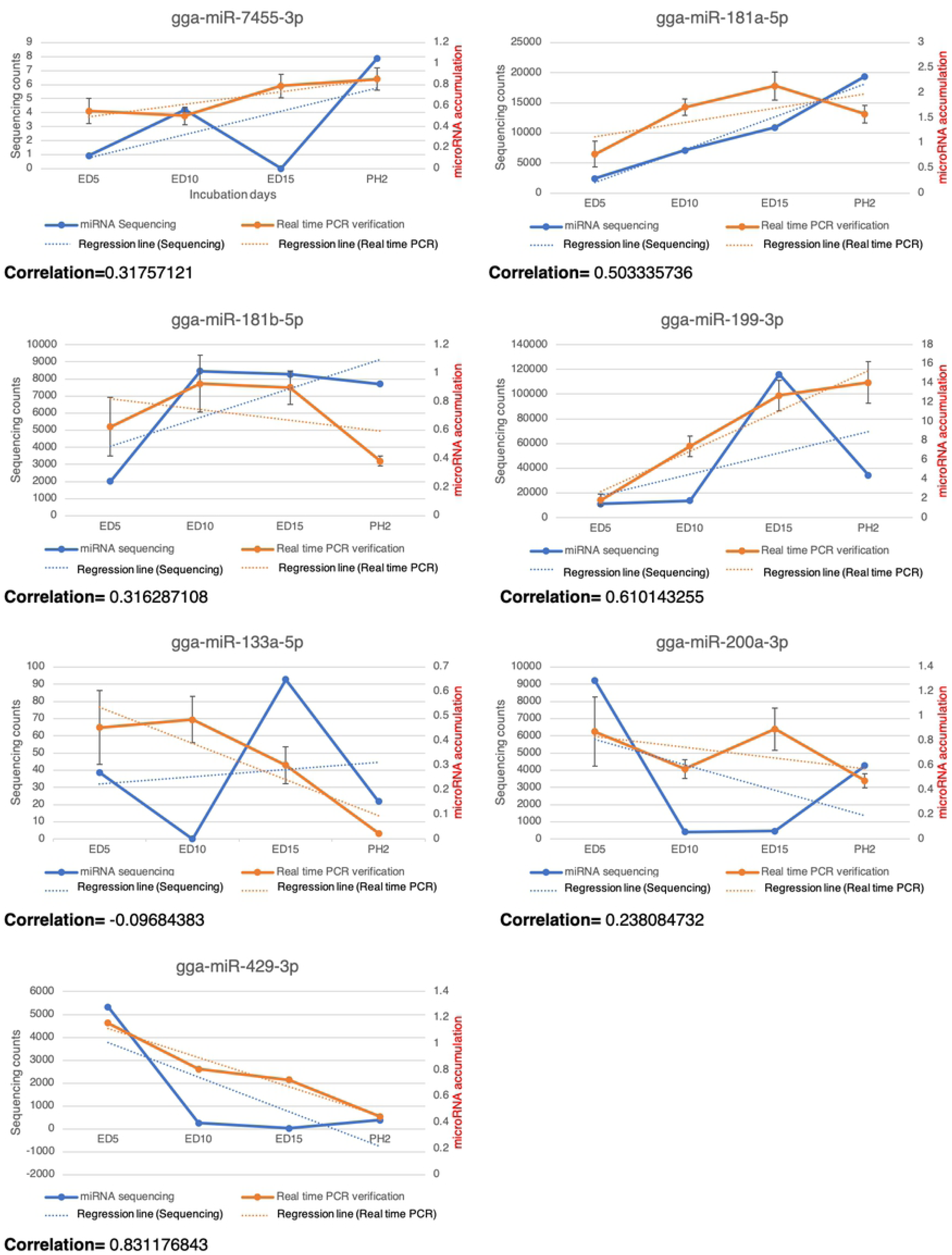
The correlations between sequencing and real-time PCR of gga-7455-3p, gga-miR-181a-5p, gga-miR-181b-5p, gga-miR-199-3p, gga-miR-133a-5p, gga-miR-200a-3p, gga-miR-429-3p. The blue lines indicated the miRNA sequencing patterns, the blue dotted lines indicated the regression lines of the sequencing data, the orange lines indicated real-time PCR results and the orange dotted lines indicated regression lines of the real-time PCR data.

### The potential functions of selected miRNAs on regulations of SOAT1 and TGFβ signaling pathway

We used the *ex vivo* culture system with EECs from Japanese quail YSMs, to study the effects of selected miRNAs. Total RNA was extracted and analyzed after transient transfection for 48 or 72 hours. The results at 48 hours showed that SOAT1 expressions were decreased by gga-miR-133a-5p and by gga-miR-429-3p; furthermore, TGFBR1 expressions were inhibited by gga-miR-133a-5p and gga-miR-429-3p. TGFBRAP1 (transforming growth factor-beta receptor associated protein 1) is a specific chaperone for Smad4 to bring Smad4 to phosphorylated Smad2/3 and to facilitate formation of the SMAD complex [18]. STRAP (serine/threonine kinase receptor associated protein) is present in a complex with Smad7 and activated TGFBR1 to stabilize the complex, and further inhibit the TGFβ signaling by preventing Smad2/Smad3 access to the receptor [19]. SMURF2 (SMAD specific E3 ubiquitin protein ligases 2) is an E3 ubiquitin ligase and can be recruited by Smad7 to form a complex to degrade TGFBR1 [20, 21].

Although the miRNAs were predicted to target genes mentioned above, expressions of TGFBRAP1, STRAP, and SMURF2 remained unchanged after transfection (Fig 3). Despite the inhibition by gga-miR-133a-5p of TGFBR1, there was no effect on SOAT1 after 72 hours transfection (data not shown). TGFBR1 is one of the receptors for the TGFβ signaling pathway. TGFBR1 is activated and phosphorylated when TGFBR2 receives ligands (e.g., TGFβ1). The downstream signals in the TGFβ signaling pathway, Smad2 and Smad3 are then phosphorylated by TGFBR1. The phosphorylated Smad2/3 joins with Smad4 to form the Smad complex and enters the nucleus for pairing with the transcription factor binding region. Furthermore, the co-repressor (e.g., TGIF) or co-activator (e.g., CBP/p300) attaches to the complex and affects the regulations of target genes.

**Fig 3.**
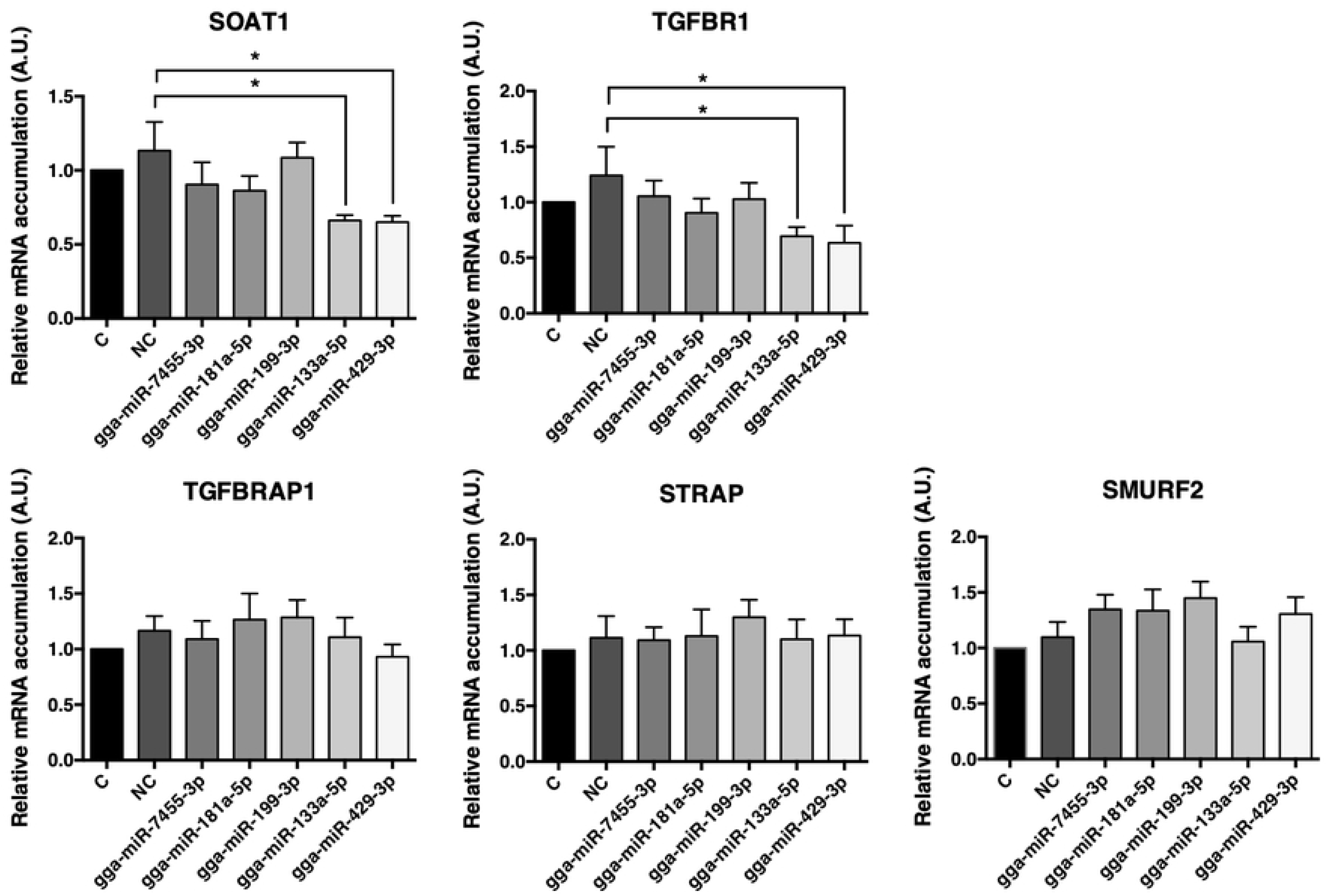
Target gene expressions after transient transfection for 48 hours using selected miRNAs. Potential target gene expressions of miRNAs transfection after 48 hours were analyzed by real-time PCR. C = control group with no transfection; NC = negative control group, the group of transfections with AllStars Negative Control siRNA. Others = groups of miRNAs transfections (5 nM miRNA mimics). N= seven to eight per group. Data were expressed as mean ± S.E.M. Control value was set as 1. All groups were compared with NC groups. Statistical significance was determined by one-way ANOVA. Dunnett’s multiple comparisons test was used to evaluate differences between means (*P ≤ 0.05).

### The validations of selected miRNAs pairing ability to the chicken TGFBR1 3’UTR

To confirm the newly-found miRNA pairing abilities to 3’UTR of the target gene, chicken TGFBR1, we constructed wild-type 3’UTR sequences of chicken TGFBR1 linked to the luciferase expression vector (Fig 4A). After co-transfection of miRNA mimics and WT-3’UTR pmirGLO plasmids into the 293T cells, the relative luciferase expressions were decreased by both gga-miR-181a-5p and gga-miR-429-3p. There was no decrease by gga-miR-133a-5p or gga-miR-199-3p (Fig 4B). The results suggested that gga-miR-181a-5p and gga-miR-429-3p target and pair with the TGFBR1 3’UTR to inhibit TGFBR1 mRNA accumulation in cells. The data also suggested that gga-miR-133a-5p may not pair with TGFBR1 3’UTR or pair outside the seed region or through other target genes to repress TGFBR1 expression in cells.

**Fig 4.**
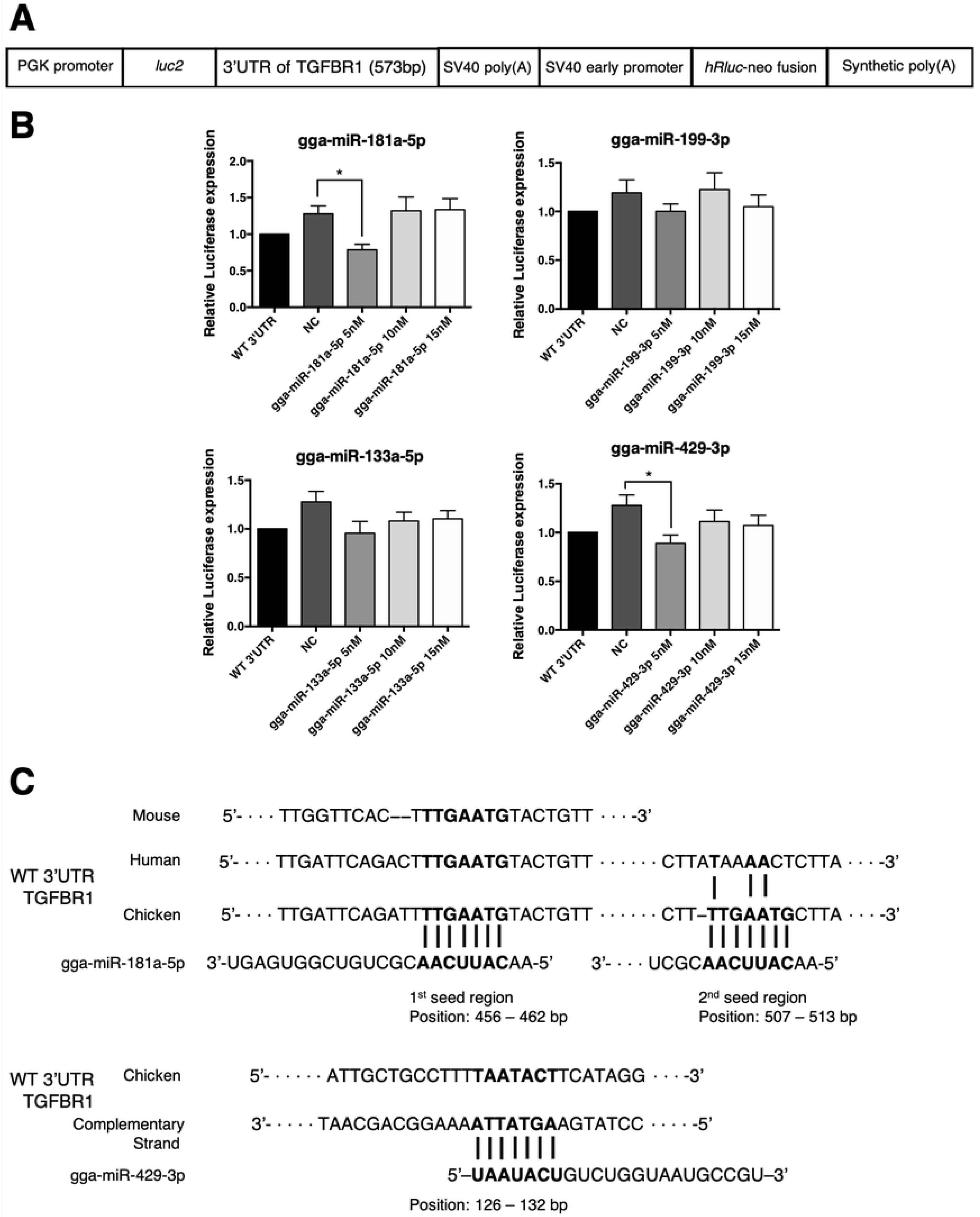
The validations of selected miRNAs to chicken TGFBR1 3’UTR. (A) The scheme of the constructed luciferase plasmid. (B) The relative luciferase expressions after miRNA mimic transfection. Transient transfection was conducted on 293T cells (3*10^4^ cells/ well). The pmirGLO-WT-3’UTR plasmid (100 ng) was used, and co-treated with 5 nM miRNA mimics or siRNA (negative control, NC). Firefly and Renilla luminescence were detected after transfection for 24 hours. N= nine per group. Data were expressed as mean ± S.E.M. Transfection with pmirGLO-WT-3’UTR only was set as 1. All groups were compared with NC groups. Statistical significance was determined by one-way ANOVA. Dunnett’s multiple comparisons test was used to evaluate differences between means (*P ≤ 0.05). (C) The sequence alignment of gga-miR-181a-5p and gga-miR-429-3p with the binding sites of the chicken TGFBR1 3’UTR.

In order to clarify the conflict between gga-miR-181a-5p transfection on gene expressions and pairing ability validation, EECs were then further transfected with different miRNA concentrations. The result showed that both SOAT1 and TGFBR1 were inhibited by 15 nM miRNA (Fig 5). Therefore, TGFBR1 expression was also regulated by gga-miR-181a-5p and gga-miR-429-3p to affect SOAT1 mRNA expression in EECs.

**Fig 5.**
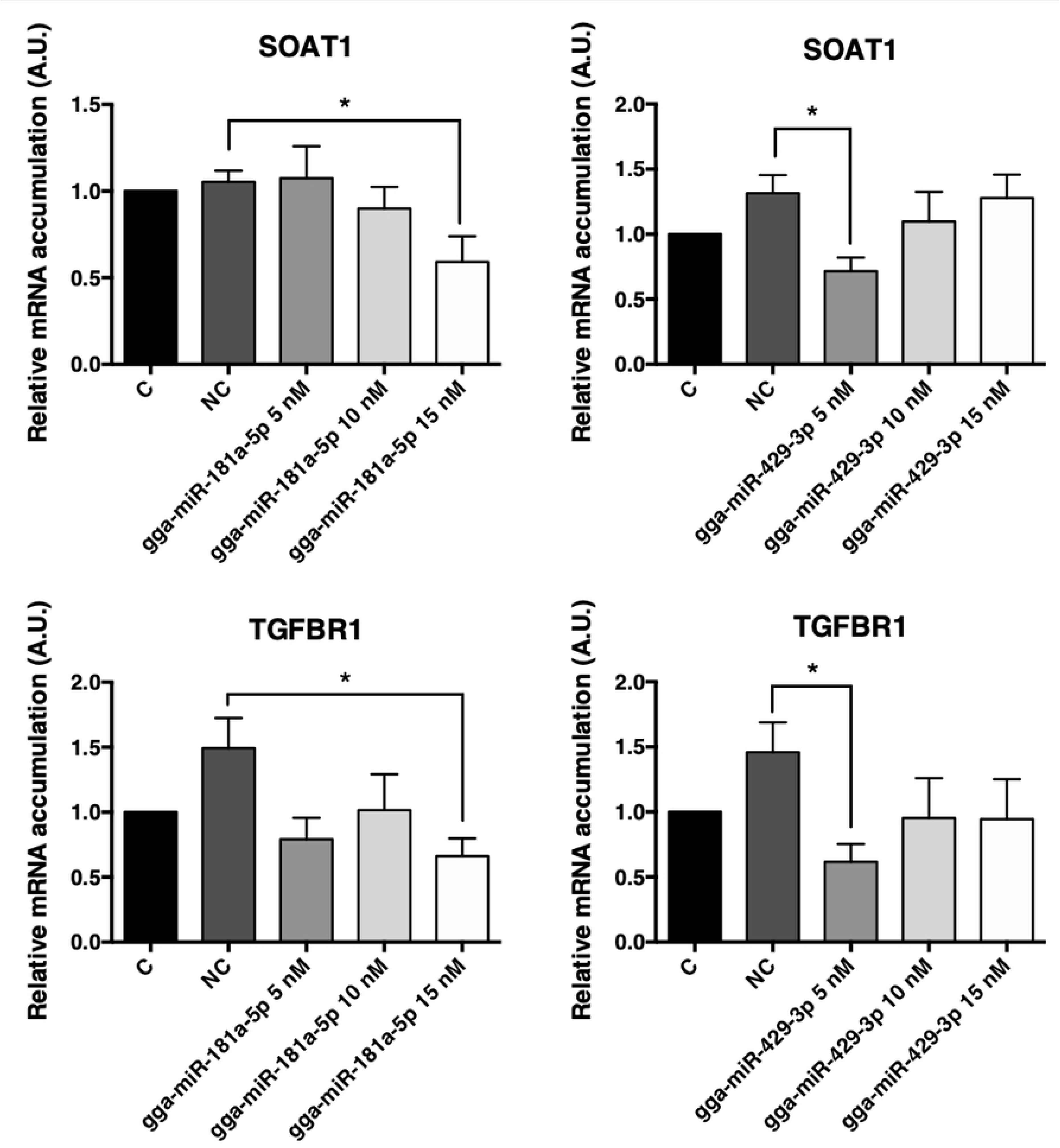
Target gene expressions after transient transfection for 48 hours using gga-miR-181a-5p or gga-miR-429-3p. The expressions of SOAT1 and TGFBR1 were analyzed by real-time PCR after 48 hours of miRNAs transfection. Data were expressed as mean ± S.E.M. N= nine per group. Control group was set as 1. All groups were compared with NC groups. Statistical significance was determined by one-way ANOVA. Dunnett’s multiple comparisons test was used to evaluate differences between means (*P≤0.05).

### Verification of interactions between selected miRNAs and the 3’UTR of TGFBR1

The miRNA pairing activities were then further compared between the WT and the mutated 3’UTR sequences of chicken TGFBR1 (Fig 6A). To determine whether the predicted seed region of gga-miR-181a-5p and gga-miR-429-3p were true binding sites, the mutated- and WT 3’UTR of TGFBR1 were separately constructed into luciferase vectors. After transfection for 48 hours, there was no difference between WT and the seeding region mutation groups with co-transfected miRNA mimics (both gga-miR-181a-5p and gga-miR-429-3p) (Fig 6B). The data revealed that gga-miR-181a-5p and gga-miR-429-3p inhibited TGFBR1 and SOAT1 mRNA expressions by directly targeting TGFBR1 3’UTR.

**Fig 6.**
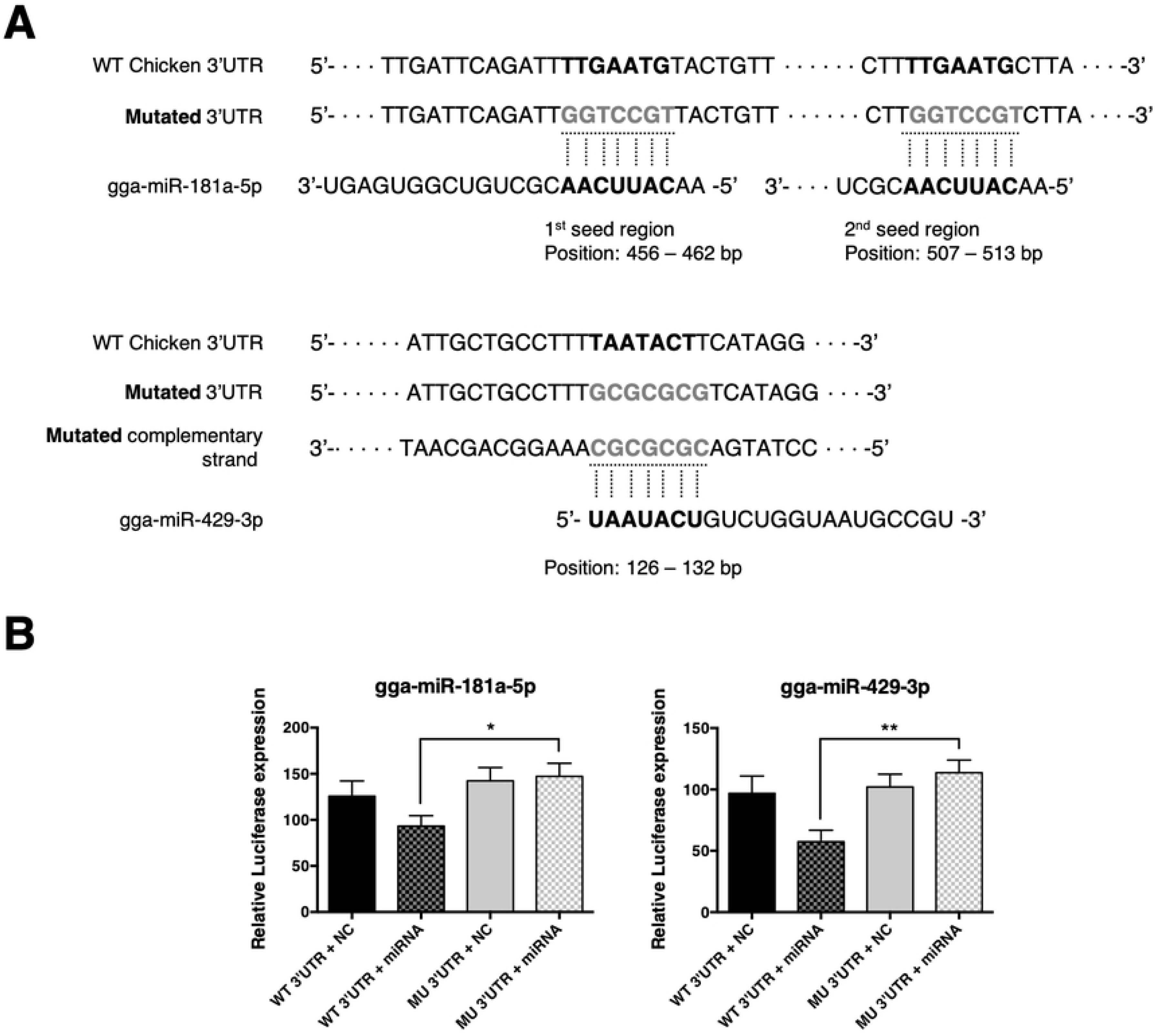
TGFBR1 is one of the direct target gene of gga-miR-181a-5p and gga-miR-429-3p. The relative luciferase expressions after miRNA mimics transfection. (A) Scheme of potential binding sites of gga-miR-181a-5p and gga-miR-429-3p on the wild- or mutated-type of chicken TGFBR1 3’UTR. (B) The transient transfection was conducted on 293T cells (3*10^4^ cells/ well). The pmirGLO-WT-3’UTR plasmid and pmirGLO-MU-3’UTR plasmid (100 ng/ well) were used and co-treated with 5 nM miRNA mimics. siRNA served as the negative control (NC). Firefly and Renilla luminescence were detected after transfection for 24 hours. N=10 to 14 per group. Data were expressed as mean ± S.E.M. All groups were compared with the NC group. Statistical significance was determined by one-way ANOVA. Dunnett’s multiple comparison test was used to evaluate differences between means. A significant difference (*P≤0.05 or **P≤0.01) was indicated.

### SOAT1 was regulated by gga-miR-181a-5p and gga-miR-429-3p by modulating TGFBR1 in the TGFβ signaling pathway

Protein concentrations of SOAT1 and TGFBR1 were examined after confirmation of two miRNAs pairing activity. Not only the mRNA accumulations were inhibited, but also both protein expression levels were found decreased after miRNAs mimic transfection for 48 hours (Fig 7), suggesting that the inhibitory effects of miRNAs were effective and consistent in EECs.

**Fig 7.**
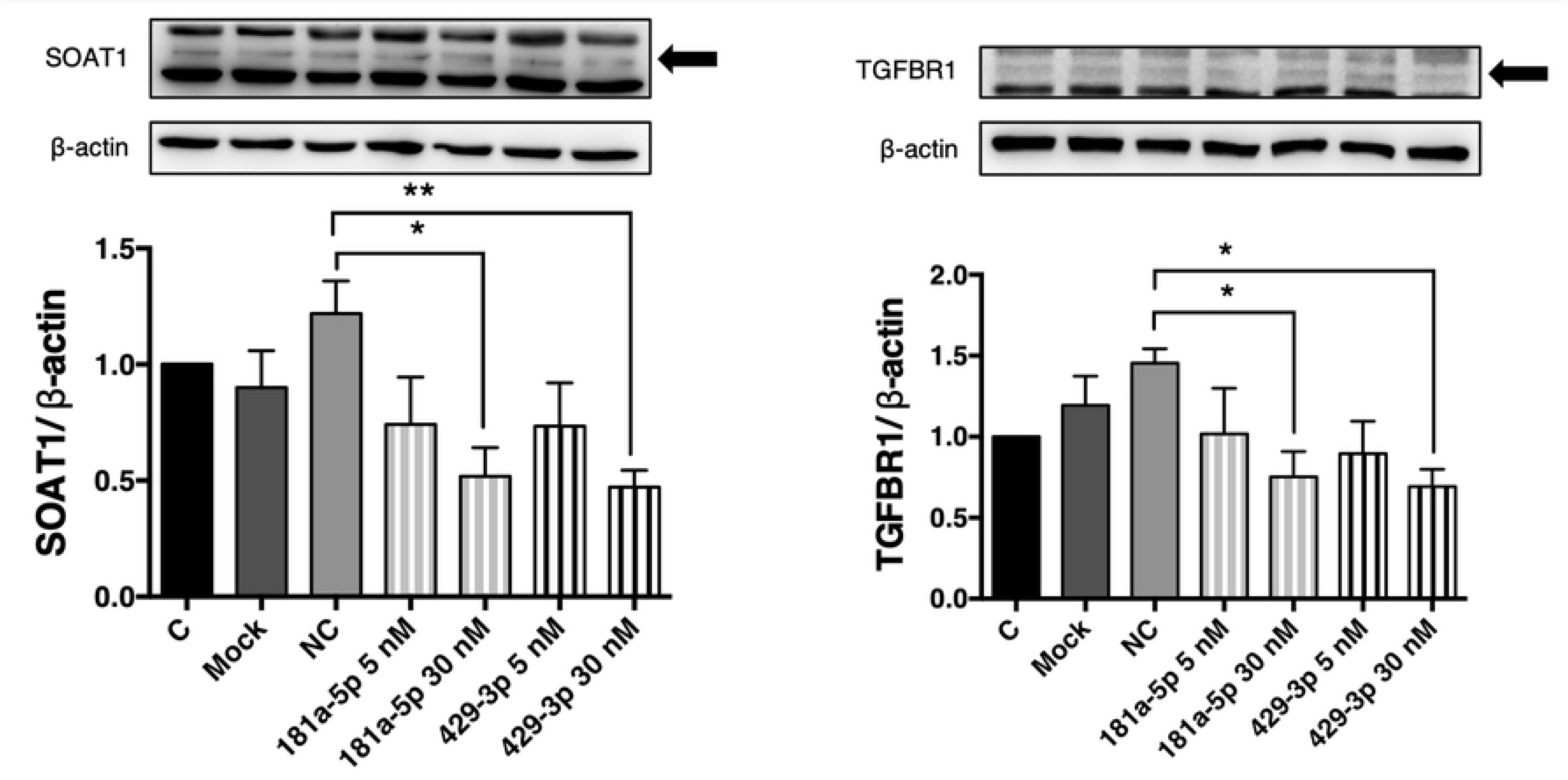
The SOAT1 and TGFBR1 protein levels after transfections with gga-miR-181a-5p or gga-miR-429-3p. EECs were transfected by miRNAs mimic for 48 hours and extracted for western blotting analysis. Total density of SOAT1 or TGFBR1 were normalized by total density of β-actin. Data were expressed as mean ± S.E.M. N = five to nine per group. C = group of no transfection in EECs, Mock = group of transfections with reagent only, NC = negative control group, the group of transfections with AllStars Negative Control siRNA. Others = groups of miRNAs transfections (5 nM or 30 nM miRNA mimics). Control group was set as 1. All groups were compared with NC groups. Statistical significance was determined by one-way ANOVA. Dunnett’s multiple comparison test was used to evaluate differences between means. A significant difference (*P≤0.05 or **P≤0.01) was indicated.

Taken together, the direct pairing ability of gga-miR-181a-5p and gga-miR-429-3p on TGFBR1 3’UTR were verified by dual-luciferase assay. The expression of TGFBR1 was directly targeted and attenuated by gga-miR-181a-5p and gga-miR-429-3p; therefore, the TGFβ pathway was affected by miRNAs, and SOAT1 mRNA and protein levels or activity was decreased. The process of cholesterol esterification was altered by miRNAs. Hence, for improving avian yolk lipid regulation to enhance hatchability during embryogenesis, it is very important to understand the involvement of miRNAs and miRNA expressions profiles in embryonic development.

## Discussions

The major findings of this study are that gga-miR-181a-5p and gga-miR-429-3p both had miRNA-mRNA interactions with TGFBR1 to produce inhibitory effects on TGFBR1 expression and regulate the TGFβ signaling pathway. In addition, the miRNAs inhibit downstream target gene expression, such as SOAT1 in EECs of Japanese quail. The pairing ability of two miRNAs to the complementary 3’UTR of chicken TGFBR1 was validated and confirmed by the dual-luciferase reporter assay. The EECs are responsible for dynamic absorption of lipids from yolk during avian embryonic development. The miRNA sequencing of YSMs revealed the miRNAs involvement during avian development. We demonstrated that SOAT1 is not only activated by a cAMP-dependent pathway [2], but also was modulated by the TGFβ signaling pathway. The current study was the first to provide direct evidence to demonstrate miRNA expression profiling in the developing YSM of avian species.

The miRNAs are highly conserved among species. Gga-miR-181a-5p shares homology with human, mouse and zebrafish, and gga-miR-429-3p is homologous with the mouse. The very first revelation of miRNA expression patterns in avian species is by whole mount in situ hybridization from the early stages, such as ED0.5 to ED5 of chicken embryogenesis [5]. This information is further expanded by miRNA sequencing for the middle stages (ED5 to ED9, and ED11) of chicken embryogenesis [22, 23]. The miRNA patterns of chicken embryonic liver or muscle of middle and later stages are also profiled and predicted to be involved in hepatocyte proliferation/lipid metabolic pathways and to regulate muscle development [6, 24]. Nonetheless, the miRNA profiling in the extraembryonic tissues such as yolk sac membranes, are less discussed during embryogenesis in avian species.

The family of miR-181a contains four members (miR-181a/b/c/d) [25]. MiR-181a-5p has been proved to have multiple functions. In dendritic cells, miR-181a-5p reduces the immunoinflammatory response from oxidized LDL in atherosclerosis by targeting the pro-inflammatory transcription factor, c-Fos [26]. In preadipocytes, miR-181a-5p induces adipogenesis by decreasing endogenous TNFα [27], or further reduces cell proliferation through the TGFβ and the Wnt signaling pathway by directly targeting Smad7 and Tcf7l2 [28]. The latest results from porcine adipose tissues indicate that miR-181a-5p directly targets TGFBR1 and enhances preadipocyte differentiation via PPARγ activation [29]. In avian species, gga-miR-181a-5p inhibits proliferation of Marek’s disease lymphoma cells by targeting MYBL1 protein [30]. High concentrations of gga-miR-181a-5p are present in the young chicken preadipocytes [31]. The circulating miR-181a-5p concentration is found low and with negative correlations in plasma triglyceride and cholesterol in hypertriglyceridemia patients. Therefore, miR-181a-5p is identified as one of the potential downregulated indicators for hypertriglyceridemia [32]. According to our results and those of prior studies, data strongly supports the involvement of gga-miR-181a-5p and the regulation of TGFBR1.

MiR-429s belong to the miR-200 family of microRNAs. MiR-429-3p has the potential to inhibit the Wnt signaling pathway and regulates adipogenesis though FABP4 activation [33]. Hsa-miR-429 inhibits epithelial–mesenchymal transition by targeting Onecut2 in colorectal carcinoma [34] and suppresses migration and invasion of a breast cancer cell line [35]. In the neurodegenerative disease aspects, levels of mmu-miR-429-3p in forebrain regions decrease in abundance at the clinical endpoint of prion disease [36]. The massive accumulation of cholesteryl ester is observed in forebrain regions from mouse models or in patients with Alzheimer’s disease (AD) [37, 38], implying that SOAT1 is actively involved in amyloid-β synthesis and AD formation. SOAT1 is one of the targets that may have beneficial effects on AD when blocked [39], and we speculate miR-429-3p may have potential relationships associated with AD.

In addition to the SOAT1 involvement in avian embryogenesis, one of the common diseases that SOAT1 may contribute to is atherosclerosis. For many years, atherosclerosis has been attributed to abnormalities in cellular cholesterol homeostasis, especially in the formation of macrophage-derived foam cells [40]. One of the cytokines known to participate in monocyte-macrophage differentiation, TGFβ1 increases SOAT1 mRNA levels in human macrophages [12]. In macrophage-derived foam cells, miR-9-5p is found to target human SOAT1 mRNA 3’UTR and to reduce SOAT1 protein levels, but not SOAT1 mRNA levels [41]. Another study shows that miR-467b directly targets mouse SOAT1 3’UTR to regulate SOAT1 and cholesteryl ester formation [42]. However, the sequences of SOAT1 3’UTR from chicken or quail are not decoded, therefore, we explored the potential upstream pathway to affect SOAT1.

According to the real-time PCR results for two different miRNAs, gga-miR-181a-5p and gga-miR-429-3p, we demonstrated that an increase in gga-miR-181a-5p levels during development of Japanese quail. In contrary to gga-miR-181a-5p levels, gga-miR-429-3p were shown decrease during the developmental process. However, we confirmed the SOAT1 and TGFBR1 inhibitions from two miRNA by EECs primary culture system. Therefore, the exact participation of gga-miR-181a-5p and gga-miR-429-3p in embryogenesis requires further examination.

The overall scheme of predicted regulation of miRNAs and the TGFβ signaling pathway with SOAT1 is illustrated in Fig 8. Taken together, embryonic SOAT1 expression in YSM was regulated by gga-miR-181a-5p and gga-miR-429-3p via the TGFβ signaling pathway, and TGFBR1 was the direct object of two miRNAs in Japanese quail. The current research found the first indication of possible regulation mechanism of avian yolk lipid utilization and modification of hatchability through changing YSM gga-miR-181a-5p and gga-miR-429-3p expressions during embryonic development.

**Fig 8.**
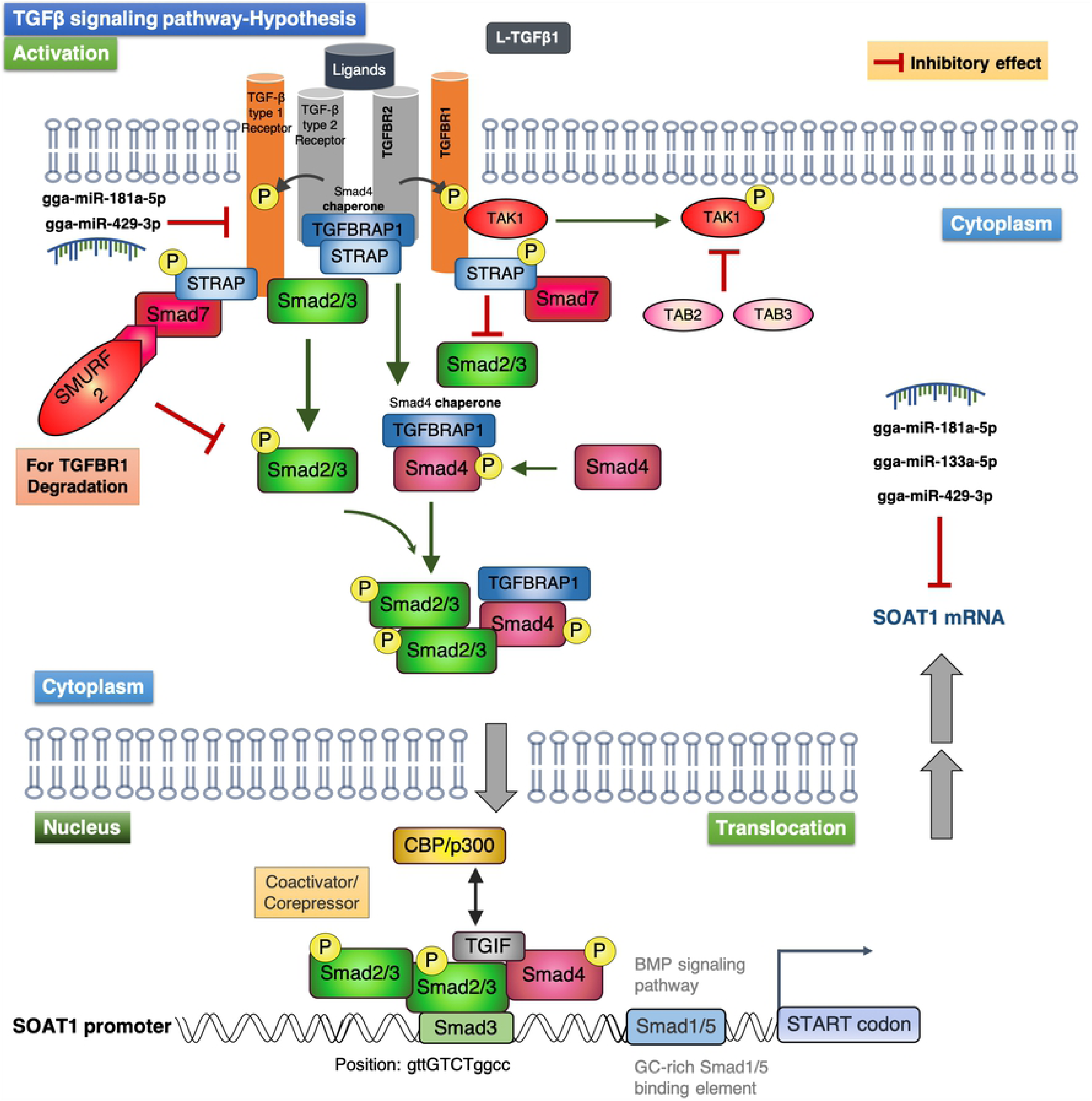
The possible relationship between miRNAs, TGFβ signaling pathway and SOAT1 expressions. The TGFβ signaling pathway is activated when the ligand (e.g., TGFβ1) binds to TGFBR2, TGFBR2 phosphorylates TGFBR1. The signal transmitter Smad2/3 is phosphorylated by TGFBR1 and joins with phosphorylated Smad4 to form SMAD complex in cytoplasm. The SMAD complex then enters nucleus to target to transcription factor binding site to affect SOAT1 gene expression. However, the gga-miR-181a-5p and gga-miR-429-3p both have the inhibitory ability on TGFBR1, and then decrease the SOAT1 expression. Although the gga-miR-133a-5p is found to attenuate SOAT1 expression, the effect is independent of the TGFβ signaling pathway.

## Conclusions

The expression profiles and involvements of miRNAs in the YSM of avian species were first demonstrated by microRNA sequencing technique. We further examined the biofunctions of gga-miR-7455-3p, gga-miR-181a-5p, gga-miR-199-3p, gga-miR-133a-5p, and gga-miR-429-3p using EECs primary culture system, and revealed the SOAT1 activity was attenuated by gga-miR-181a-5p and gga-miR-429-3p through directly inhibiting TGFBR1 in the TGFβ signaling pathway. This was indicative of possible regulations of avian yolk lipid utilization and modification of hatchability by changing predicted miRNA expressions.

## Acknowledgements

This work was supported by the Ministry of Science and Technology, under Grant MOST 107-2313- B-002-050-MY3 in Taiwan. We especially thank the Technology Commons, College of Life Science, National Taiwan University for technical assistance on using the SpectraMax i3x Multi-Mode Plate Reader and the ultracentrifugation. We declare that the experiments comply with the current Taiwan laws, the place in which the experiments were performed. There is no conflict of interest.

